# Patterns of low-complexity regions in human genes

**DOI:** 10.1101/2023.12.01.569686

**Authors:** Lokdeep Teekas, Nagarjun Vijay

**Author notes:** Correspondence: Nagarjun Vijay, **Email:**. **Author Contributions:** NV conceived the idea. LT and NV designed the analyses and wrote the manuscript. **Competing Interest Statement:** Authors declare no competing interests.

## Abstract

Genome evolution stands as a paramount determinant for species survival and overall biodiversity on Earth. Among the myriad processes orchestrating genome evolution, the dynamic attributes of length and compositional polymorphism within low-complexity regions (LCR) are the fastest. Clusters of LCR hotspots serve as pivotal conduits connecting different modes of genome evolution, specifically arising through gene duplication events and harboring pivotal sites susceptible to point mutations. Thus, they offer a holistic perspective on the panorama of genome evolution. Furthermore, LCR actively participates in a multifaceted spectrum of neurological, developmental, and cognitive disorders. Despite the substantial body of knowledge concerning the roles of individual LCR-containing genes in the causation of diseases, a comprehensive framework remains conspicuously absent, failing to provide a unified portrayal of LCR-containing genes and their interactions. Furthermore, our understanding of the intricate interplay between paralogy and LCR remains notably deficient. Within this study, we have identified nine clusters of LCR hotspots within the human genome. These clusters are predominantly comprised of closely positioned paralogs, characterized by a significantly higher prevalence of shared LCR and a lower degree of differentiation (F_ST_) across diverse human populations. Moreover, we have unveiled intricate networks of LCR-containing genes engaged in mutual interactions, sharing associations with a spectrum of diseases and disorders, with a particular emphasis on hereditary cancer-predisposing syndromes. Our discoveries shed light on the compelling potential of LCR-containing interacting genes to collectively engender identical diseases or disorders, thereby underscoring their pivotal role in the manifestation of pathological conditions.

**Significance Statement:** Among myriad genome evolution processes, low-complexity regions (LCR) are pivotal, being both the fastest and bridging other evolution modes like gene duplication and point mutations. Understanding LCR-containing paralogous genes is essential to comprehend genetic diseases. Here, we demonstrate that the human genome harbors clusters of LCR hotspots mainly composed of paralogous genes sharing LCR, indicating a role for segmental duplication. The degree of differentiation is significantly lower in clusters of LCR hotspots than in other regions. Moreover, we provide a detailed network of LCR-containing interacting genes associated with shared diseases. Instead of attributing a single disease to an LCR gene, a unified perspective on LCR-containing interacting genes causing the same disease enhances our understanding of LCR-induced disease mechanisms.

## Introduction

Genome evolution is one of the most fundamental aspects of a species’ survival and overall biodiversity on Earth. It is well-established that genome evolution is governed by various mechanisms, including point mutations, gene duplications/losses, segmental insertions/deletions, rearrangements, and horizontal gene transfers (1–3). However, another prominent phenomenon, exhibiting rates up to 100,000 times higher than point mutations, is the expansion and contractions of low-complexity regions (LCR) (4–6). In the amino acid sequence context, the regions of amino acids (single or multiple) characterized by a reduced diversity of residues below the average unbiased composition are termed low-complexity regions (LCR) (7–9). While LCR within coding regions were previously conjectured to be analogous to “junk” DNA sequences, it is now firmly established that they play pivotal functions in numerous essential biological processes, including recombination (10), protein kinases (11), transcription regulation (12, 13), as well as neurogenesis, DNA repair (10), cell adhesion, development, cognition, and immunological responses (14–16), among others. Specific LCR, such as polyL stretches, are recognized for their dual utilization by hosts and pathogens in host-pathogen interactions. Pathogens employ leucine-rich repeats (LRRs) to adhere to host cells and gain entry (14). Conversely, host cells employ LRRs to recognize specific pathogen-associated molecules and activate the innate immune system (14, 17). In addition to their indispensable role in biochemical functions and immunological responses, LCR also plays a pivotal role in morphology and physiology, facilitating rapid adaptation and evolutionary novelty, for which they are famously termed ‘tuning knobs’ (18). LCR serves as a platform for selection by generating rampant mutations that contribute to functional and morphological diversification (6, 19), with certain amino acid tracts quantitatively linked to an organism’s phenotypic traits, as exemplified by the polyQ/polyA ratio in RUNX2 associated with clinorhynchy (6, 20). Furthermore, LCR play a crucial role in numerous neurodegenerative diseases, cognitive disabilities, and cancers (10, 21–25). One of the most compelling illustrations of LCR involvement in diseases is Huntington’s disease. The *HTT* gene encodes the huntingtin protein, which plays a significant role in brain neuronal development during the embryonic stage (26, 27). Exon 1 of the *HTT* gene harbors a CAG repeat, resulting in a PolyQ stretch. This region comprises 8 to 35 Q residues in the wild-type huntingtin protein, while patients with Huntington’s disease exhibit 36 to 180 residues (28).

Processes such as replication slippage, unequal crossing-over, and gene conversion (recombinational mechanisms) can give rise to both the origin and length variability of LCR (4, 11). Newly emerged LCR commonly undergo periods of positive or relaxed purifying selection before attaining their selection equilibria (29). Depending on their functional requirements and the selective pressures they encounter, orthologous LCR can exhibit either variability in length and composition or remain highly conserved across species. The volatility in the length of LCR and their intrinsically unstable nature enables them as one of the most prominent phenomena for genome evolution.

Among other mechanisms for genome evolution, gene duplication is of immense importance. The newly emerged gene copies are often conditionally exempted from selection pressure, providing an opportunity for neo- or sub-functionalization (30, 31). Segmental duplication constitutes the primary mechanism for the generation of proximal paralogous genes, thereby providing opportunities for adaptive evolution through gene fusion and exon exaptation (32, 33). Gene paralogy not only plays a prominent role in evolution, but is also a leading cause of many genetic defects, especially which are dosage-dependent during embryonic development (34, 35).

While both gene paralogy and LCR offer avenues for genomic innovations and contribute to genetic diseases, their interaction remains understudied (3). Genes containing LCR, when duplicated, have the potential to increase the LCR repertoire within the genome. This augmentation in LCR content may promote genomic innovations but also elevate the likelihood of genetic diseases and disorders. Understanding the interplay of gene paralogy and LCR is imperative for comprehending development, disorders, and evolution, especially in the context of LCR-containing paralogous genes. To investigate the evolutionary implications of LCR-containing paralogous genes in genetic diseases and disorders, we have chosen the human genome as our study focus due to its comprehensive database of mutations leading to genetic diseases and disorders (36). Despite the wealth of knowledge regarding the involvement of LCR in diseases, an integrated perspective on the collaborative influence of LCR-containing genes in causing the same disease remains unexplored. In this study, we comprehensively analyzed all annotated protein-coding genes within the human genome, highlighting clusters of LCR hotspots. These clusters harbor LCR-containing paralogous genes in close proximity, suggesting a role for segmental duplication. Furthermore, our findings indicate that LCR-containing interacting genes have the potential to engender the same disease or disorder.

## Results

### DNA repeats may contribute to amino acid LCR

Utilizing data on DNA repeat distribution within the human genome from the UCSC genome browser, we conducted an investigation into the distribution of these repeats, specifically within exonic regions. Our analysis revealed a pronounced abundance of DNA repeats within human exons, with short interspersed elements (SINE) exhibiting the highest frequency, followed by simple repeats, long interspersed nuclear elements (LINE), and DNA repeats (Fig. S1). Upon further examination of the distribution of amino acid LCR, we observed that proline-rich regions predominate within human exons, followed by isoleucine (Fig. S2). Glutamate, serine, and leucine LCR displayed similar levels of abundance within the exonic regions.

Previous studies have indicated that stretches of high GC content can facilitate the formation of LCR (37, 38). Our observations corroborate this, as exons containing LCR demonstrated GC content levels between 35% and 55% (Fig. S3), whereas exons lacking LCR occasionally exhibited GC content exceeding 60%. Moreover, we explored the potential for DNA repeats within coding regions to give rise to amino acid LCR. Interestingly, while SINE represents the most prevalent DNA repeat in the human genome, our analysis revealed that simple DNA repeats primarily contribute to the formation of amino acid LCR (Fig. S4), followed by low-complexity DNA repeats and SINE. These findings shed light on the distinctive roles of different DNA repeats in shaping the landscape of amino acid LCR within coding regions.

### The human genome contains hotspots of LCR

The designation “LCR hotspot” in the genome is conferred upon the top 1% region within a 50Kb window, determined based on the ratio of LCR fraction to exon fraction. Furthermore, we define a “cluster of LCR hotspots” as a region spanning 10Mb and containing more than five LCR hotspots. In our investigation, we have identified nine such clusters of LCR hotspots within the human genome (Fig. 1a). Examining the contributions of various DNA repeats to amino acid LCR hotspots, we find that simple repeats, low-complexity sequences, LINE, and SINE elements are the primary contributors. Notably, while SINE elements are more abundant than LINE elements in the broader context of LCR, our analysis reveals a shift in abundance within LCR hotspots (Fig. S5).

**Figure 1.**
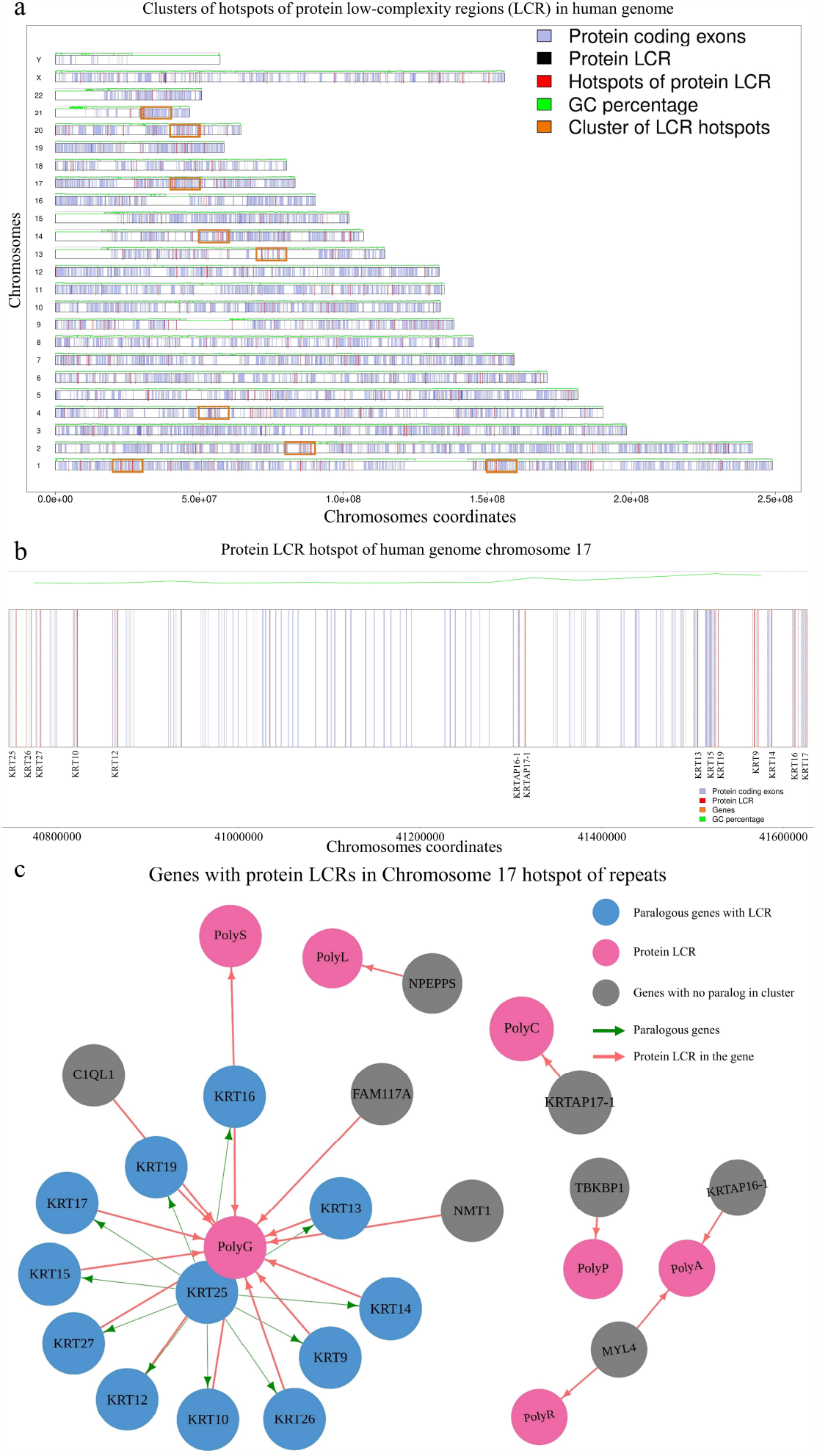
The human genome contains clusters of LCR hotspots comprised of paralogous genes. (A) The distribution of LCR hotspots is shown with vertical red dashed lines, while clusters of LCR hotspots are enclosed in orange boxes within protein-coding genes. Exons are depicted by faded-blue bars. The green line above each chromosome represents the GC content in a 50 Kb non-sliding window, with a black dashed line denoting the 50% mark. (B) A specific cluster of LCR hotspots on chromosome 17 highlights the presence of keratin paralogous genes. The orange line represents the gene’s length, and faded-blue bars indicate exons. LCRs within the exons are marked with red bars. Genes without LCRs are not labeled for clarity. (C) The cluster of hotspot LCRs on chromosome 17 primarily consists of paralogous genes, with genes from the same family represented in the same color (in this case, blue). Genes without paralogs within the cluster are depicted in gray. LCRs are indicated with the hotpink2 color, while indianred1-colored arrows denote the presence of LCRs within the genes, and green4 represents the paralogous genes.

To gain deeper insights into these clusters of LCR hotspots, we have selected two specific clusters, one located on chromosome 17 (Fig. 1b) and another on chromosome 1. Our observations indicate that both hotspots are predominantly comprised of closely spaced paralogous genes. The hotspot on chromosome 17 primarily features genes from the keratin family, with a notable prevalence of polyG LCR (Fig. 1c). Conversely, the hotspot on chromosome 1 consists mainly of late cornified envelope proteins sharing a polyQ region (Fig. S6). These findings offer valuable insights into the composition and characteristics of LCR hotspots in specific genomic regions.

### Paralogous genes contribute to LCR hotspot

To assess the contribution of LCR-containing paralogous genes situated in close proximity within hotspots, we conducted a comparative analysis between the number of paralogs within hotspots and those in non-hotspot regions, as well as the proportion of shared LCR (common LCR) among paralogs within hotspots versus those in non-hotspot regions. Our findings reveal that hotspots exhibit a significantly greater abundance of paralogs (Wilcoxon test; p < 0.05) (Fig. 2a) and paralogs sharing common LCR (Fig. 2b) within the human genome. Notably, a high mutation rate or the presence of polymorphic sites can result in reduced levels of population differentiation, as indicated by the fixation index (F_ST_). In cases where a locus exhibits polymorphism, it can yield a low F_ST_ value, even in the absence of shared alleles between different populations (39–41). Given that LCR encompasses rapidly evolving sites, it was anticipated that LCR would exhibit a low degree of population differentiation, as indicated by F_ST_. A pairwise comparison of F_ST_ values among five major human populations indeed unveiled a notably reduced level of differentiation in LCR compared to non-LCR regions (Wilcoxon test; p < 0.05) (Fig. 2c). Furthermore, an examination of the folded site frequency spectrum of minor allele frequencies for exons harboring LCR and those without LCR revealed a comparable pattern (Fig. S7).

**Figure 2.**
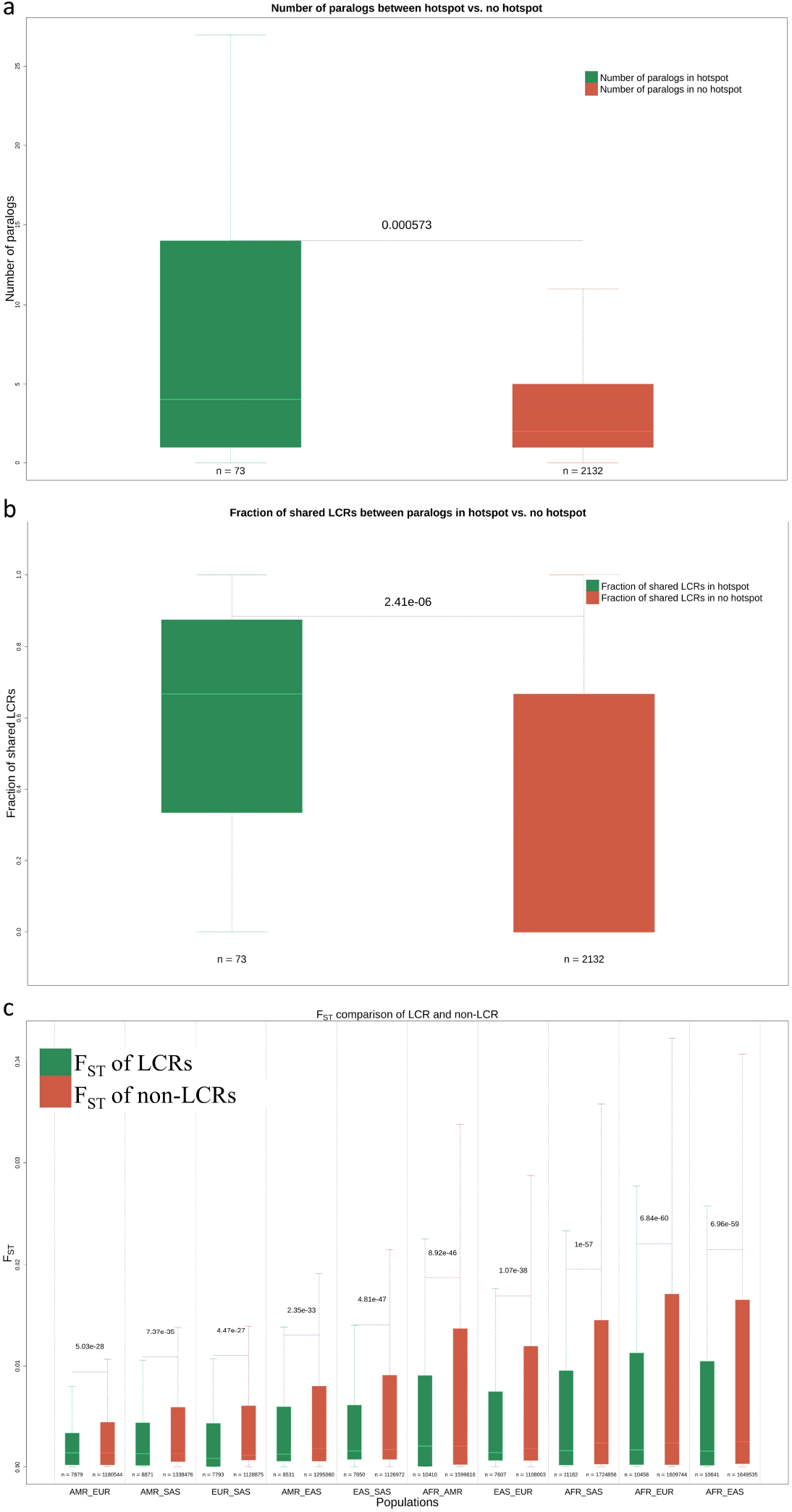
The LCR hotspots exhibit a notably higher number of paralogs that share LCR sequences. (A) Within the human genome, LCR hotspots contain a significantly greater number of paralogous genes (73 genes in hotspots versus 2132 genes in non-hotspot regions) compared to regions lacking hotspots (Wilcoxon test, p < 0.05). (B) Genes within LCR hotspots also display a significantly increased number of shared LCR sequences (the same LCR sequences within genes) compared to genes in non-hotspot regions (Wilcoxon test, p < 0.05). (C) Furthermore, the degree of differentiation (FST) in LCR sequences is significantly lower (Wilcoxon test, p < 0.05) in all pairwise comparisons among major human populations.

### Role of LCR-containing genes in disorders

LCR play a pivotal role in a diverse spectrum of neurological, developmental, and psychological disorders and diseases. While a considerable body of knowledge underscores the involvement of gene-specific LCR in these conditions, a holistic perspective on interacting LCR-containing genes remains relatively scarce. In pursuit of a comprehensive understanding, we investigated the role of interacting genes that share at least one LCR and contribute to the same disorders and diseases. Our initial exploration revealed that genes featuring polyA, polyE, polyG, and polyL regions are disproportionately involved in disorders compared to their counterparts (Fig. S8).

Furthermore, among the 1154 LCR-containing interacting genes associated with disorders, we observed that 490 genes harbor at least one paralog within this list, constituting 42.46% of LCR-containing genes that have LCR-containing paralogs and are linked to disorders. Additionally, within the cohort of 791 instances of disorders featuring LCR-containing paralogs, 55 pairs exhibit shared LCR and are concurrently associated with the same disorder (Fig. S9). Our observations further unveiled the presence of multiple communities of LCR-containing interacting genes across the genome, collectively contributing to shared diseases and disorders (Fig. 3), with a particular emphasis on hereditary cancer-predisposing syndromes (Fig. S10). This comprehensive exploration sheds light on the intricate web of interactions among LCR-containing genes in the context of diseases.

**Figure 3.**
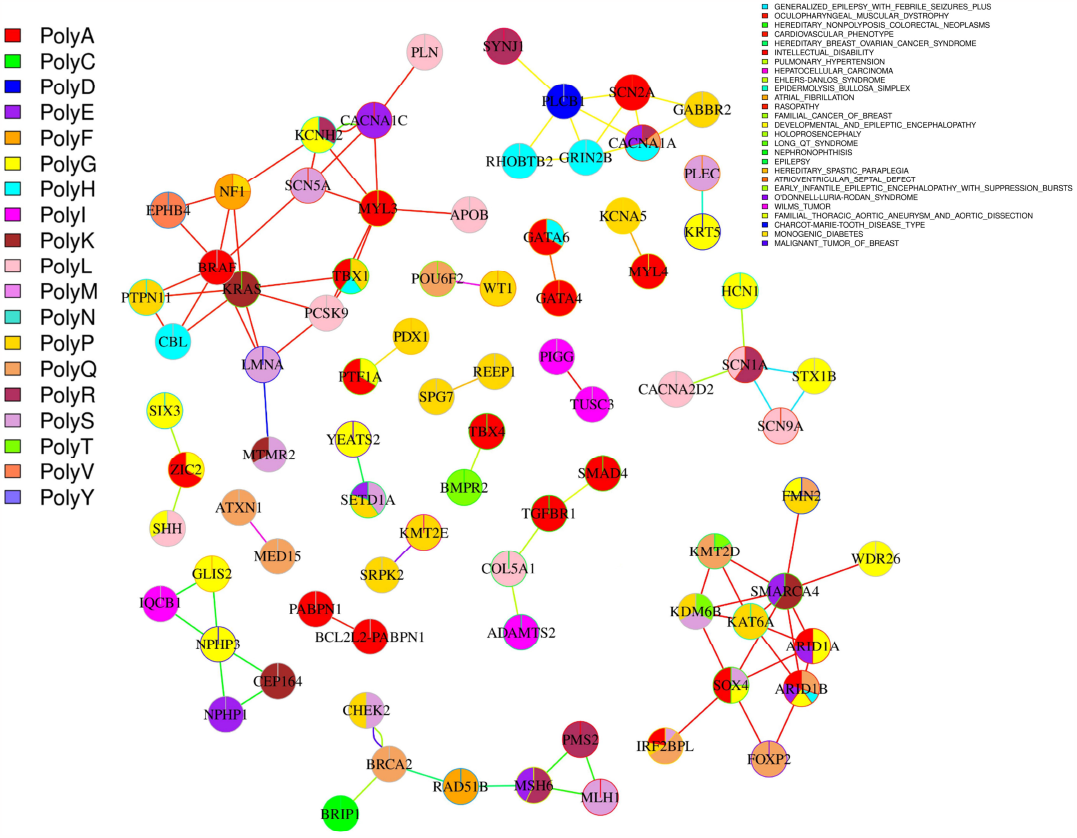
This network comprises genes that contain low-complexity regions (LCR) and are involved in causing diseases through their interactions. We have chosen to display only experimentally validated interactions where genes with LCR contribute to the development of at least one common disease or disorder. Within each vertex, a pie chart illustrates the proportion of LCR within the gene. The vertex frame indicates paralogous genes, while a gray color signifies the absence of paralogs. Shared diseases are depicted by edges of the same color within the network.

## Discussion

Genome evolution holds critical significance by serving as the supporting framework of biodiversity, explicating the genetic basis of phenotypic traits and diseases, guiding conservation strategies, strengthening technological advancements, and providing insights into the life histories on Earth. Consequently, the investigation of genome evolution carries profound implications across the diverse realms of biology, agriculture, environmental science, and medicine. LCR and gene paralogy represent two crucial mechanisms in genome evolution, making it intriguing to explore their interplay. LCR contribute to rapid evolution by facilitating molecular innovation not only through length variation, which can influence protein structure, but also by harboring sites for point mutations. Gene paralogy, on the other hand, augments the gene repertoire within the genome, thereby fostering opportunities for evolutionary novelty.

In our initial analyses focused on elucidating the contribution of DNA repeats to the formation of amino acid LCR (later referred to as LCR for amino acid LCR), we observed a notable prevalence of DNA repeats within LCR. This observation aligns with expectations, as repetitive DNA regions within coding sequences often give rise to repetitive amino acid sequences. However, it is worth noting that a region containing an LCR may not be identified as harboring DNA repeats due to codon degeneracy. The elevated abundance of DNA repeats within LCR suggests the potential influence of replication slippage in their generation, or alternatively, insufficient time may have elapsed for point mutations to degrade or degenerate the sequence. Similar to a previous study, we also observed a high abundance of proline, isoleucine, glutamic acid, and serine LCR in the human genome (42). Intriguingly, proline stretches exhibit the lowest codon homogeneity (CH: length of the longest pure codon stretch/size of the repeat region), with a CH value of 0.4, along with the smallest mean length of the longest pure codon stretches and the lowest percentage of LCR encoded by pure codons (3%) in both human and mouse genomes (43). A relatively higher abundance of simple DNA repeats than any other repeat type suggests a prominent role of replication slippage in the origin of LCR. However, the persistence of such a high abundance of proline LCR alongside very low CH values suggests that replication slippage alone cannot account for these patterns and implies a crucial role for selection in shaping these characteristics.

An examination of LCR within genes has elucidated the presence of clusters of LCR hotspots within the human genome. These regions are characterized by an excess of LCR-containing genes, with a substantial proportion of LCR sequences residing within these genes. It is intriguing that these clusters exhibit a significantly higher number of paralogs in close spatial proximity, as well as shared LCR between the paralogous genes. This implies the possibility of the recent emergence of these paralogs through segmental duplications within an evolutionary context or the potential coferment of advantageous features by LCR, thereby providing opportunities for functional innovations. Especially, the presence of glycine-rich regions in keratin family genes is intriguing. Glycine-rich regions are known to form interaction sites of various proteins and participate in prion-protein interaction in humans (44, 45). In plants, glycine-rich proteins provide adaptive abilities by participating in stress responses like wound healing, viral infection, exposure to cold, and water stress (46, 47). The keratins represent a highly diverse family of intermediate filament proteins, with their presence primarily organized into two clusters, notably type ? on chromosome 17 and type ? on chromosome 12, within the human genome (48–50). Keratin proteins participate in a variety of functions in the cytoskeletal system, epithelial tissues, and morphological features and provide mechanical stability to tissues (48, 51). Recent investigations have underscored the pivotal role of keratins in numerous hereditary diseases as well as in the context of various cancers (52–54). The presence of glycine-rich regions within keratins suggests a potential involvement of LCR in enhancing the interacting capabilities of these proteins in various functional contexts.

Although the site frequency spectrum of minor allele frequencies in LCR and non-LCR regions displayed no discernible distinctions, in a compelling pairwise assessment of the degree of differentiation among major human populations (AFR, AMR, EUR, EAS, and SAS), the F_ST_ (Fixation index) of LCR is significantly higher compared to regions without LCR. LCR are well-documented for their propensity to facilitate rapid diversification through both length and point mutations (12, 18). The substantial elevation in F_ST_ within LCR implies that the genetic sites encompassed by LCR exhibit ongoing and rapid variation, thereby limiting the opportunities for natural selection to fix a specific allele within the population.

LCR confer not only substantial evolutionary advantages but also serve as causal factors in a spectrum of genetic diseases and disorders (55–57). Isolated investigations of individual disease-associated genes offer limited insights into the broader evolutionary implications of LCR involvement in such conditions. Therefore, it is imperative to establish a comprehensive perspective on LCR-containing genes that contribute to shared diseases or disorders. Our analysis has unveiled clusters of interacting genes that, despite not being paralogs, share LCR sequences and are associated with identical diseases or disorders. In certain instances, the interaction between two LCR-containing genes can give rise to multiple shared diseases. Notably, LCR have recently gained recognition as contributors to genomic instability leading to tumor progression and the development of cancers (58). We have identified a specific cluster of interacting LCR-containing genes that are also linked to hereditary cancer-predisposing syndromes. The presence of a substantial number of LCR-containing interacting genes organized into clusters associated with hereditary disorders underscores their participation as a coordinated network rather than functioning as individual, isolated genes. This networked involvement suggests a complex interplay among these genes, collectively contributing to the manifestation of hereditary disorders instead of any single gene in isolation.

In conclusion, LCR are the “double-edged sword” of genome evolution. Particularly, LCR-containing paralogous genes congregated within LCR hotspots predominantly originate from segmental duplications, which stand as another prominent mechanism in the framework of genome evolution. These LCR-containing genes further accommodate sites prone to point mutations, a substantial mechanism that expedites rapid adaptation. Consequently, LCR function as pivotal connectors bridging diverse modes of genome evolution. In addition to their role as facilitators of genome evolution, LCR concurrently emerge as prominent causal agents underlying a spectrum of hereditary, developmental, neurological, and cognitive disorders. A comprehensive examination of LCR-containing interacting genes, which contribute to shared diseases, shines a spotlight on their hitherto underappreciated involvement in both cancers and hereditary disorders. This multifaceted role of LCR underscores their significance in shaping the genetic landscape and warrants further exploration in understanding complex disease mechanisms.

## Materials and Methods

### Dataset preparation

In our study, we downloaded the human protein-coding genes’ coding sequences and their translated sequences from Gencode Release 42 (GRCh38.p13) (59). We utilized *CompositionMaker* to compute the background frequency of each amino acid residue in the translated sequences and detected low-complexity regions (LCR) using fLPS2.0 (60). The detected LCR were then filtered based on the following criteria:

- LCR should be longer than three amino acids.
- The region’s base composition should not contain more than four unique amino acid residues.
- The region should not include an “X” in its composition.
- The purity of the region, defined as the number of residues of the base composition divided by the total number of residues in the stretch, had to be greater than fifty percent.

We then mapped the filtered LCR to their corresponding genomic coordinates using PoGo (61) and eliminated regions with multiple mappings. The data about DNA repeats were obtained from the UCSC web server, encompassing exonic and non-exonic repeats. By utilizing bedtools (62), we intersected the DNA repeat file with the amino acid LCR file to ascertain the involvement of DNA repeats in the formation of amino acid repeats. Additionally, we calculated the %GC content in the human genome using bedtools (62) in 50kb non-overlapping windows.

### Identification of hotspots of protein LCR

To identify amino acid LCR hotspots within the genome, we determined the fraction of bases covered by exons and LCR within each 50kb window. The LCR fraction was divided by the exon fraction, and we identified the top 1% fraction as indicative of amino acid repeat hotspots. We counted the number of repeat hotspots in 1Mb non-overlapping windows using bedtools to identify clusters of repeat hotspots. A window containing more than five repeat hotspots is considered a cluster of hotspots. We identified 73 genes present in the cluster of hotspots.

From the clusters of LCR hotspots, we selected two clusters for better visualization: one from chromosome 1 and one from chromosome 17. Furthermore, for the genes present in the cluster, we identified their paralogs, using Ensembl (63), within the same cluster and their LCR. We generated gene networks for the genes based on their paralogs and shared LCR using the igraph (64) package in R.

### Distribution of DNA repeats and amino acid LCR

To quantify the occurrence of each amino acid LCR in the human genome, we selected the output of fLPS2.0 and counted each LCR type using shell scripting. We quantified the abundance of DNA repeats in the exonic regions by intersecting the UCSC-detected DNA repeat coordinates with the exon coordinates using bedtools intersect and counting each DNA repeat type using shell scripting. To get an insight into the abundance of DNA repeats in the amino acid LCR in the hotspots, we intersected the coordinates of DNA repeats with the coordinates of the LCR in the hotspot regions. Using R, we also calculated the relative abundance (proportion) of different DNA repeats in amino acid LCR.

### Population genetics of LCR

We downloaded the VCF files for each chromosome with the extension “filtered.SNV_INDEL_SV_phased_panel.vcf.gz” from the 1000 Genomes EMBL-EBI ftp server, concatenated all the VCF files, intersected the VCF file with LCR-containing exons and no LCR exon coordinates using bedtools, calculated the minor allele frequency (MAF) using VCFtools (65), and visualized the folded site-frequency spectrum (SFS) using R.

We subsetted the concatenated VCF file according to the populations (AMR, AFR, EUR, EAS, and SAS) and calculated F_ST_ using the “--weir-fst-pop” argument from the VCFtools for all pairwise combinations of populations. The output is subsetted according to overlap with LCR. We compared the distribution of F_ST_ of LCR with non-LCR using the Wilcoxon test (p < 0.05 for significance) for each pair of populations.

### Disease network of LCR-containing genes

We downloaded the pairwise list of all the experimentally validated interacting proteins from the STRING database v12.0 (66). We downloaded the information about genetic diseases and disorders from the ClinVar (36) database. Using bedtools intersect, we selected the disorder-causing variations that overlap with LCR. We generated a network of the experimentally validated interacting genes using the igraph package in R. We color-coded the edges, vertices, and vertex frame based on the following criteria:

- The pie chart inside each vertex represents the gene’s proportion of amino acid LCR.
- The frame/boundary color of the vertex represents whether the gene has a paralog within the network. Paralogs have the same vertex frame color, while grey means no paralog in the network.
- The edge color signifies the shared disease/disorder between two interacting genes.

To gain better visual insight, we constructed a gene-interacting network of the hereditary cancer-predisposing syndrome separately.

### Statistical analyses

All the necessary statistical analyses and visualization are performed in R. The results with p < 0.05 are considered statistically significant.

## Data and code availability

All the necessary data and codes to generate the results are available in the Github repository: https://github.com/ceglablokdeep/human_genome_LCRs

## Acknowledgments

We thank the Ministry of Human Resource Development for fellowship to LT. Computational analyses were done on the Har Gobind Khorana Computational Biology cluster established and maintained by combining funds from IISER Bhopal under Grant # INST/BIO/2017/019, IYBA 2018 from the Department of Biotechnology (Grant no. BT/11/IYBA/2018/03), and ECRA from Science and Engineering Research Board (Grant no. ECR/2017/001430).

