## Supplementary Text and Figures for "Patterns of low-complexity regions in human genes"

**This file includes:**

Figs. S1-S6. The figures S7-S10 are uploaded separately. Legends are provided here.

**Data and code availability:**

All the necessary data and codes to generate the results are available in the Github repository: github.com/ceglablokdeep/human_genome_LCRs

### Figures

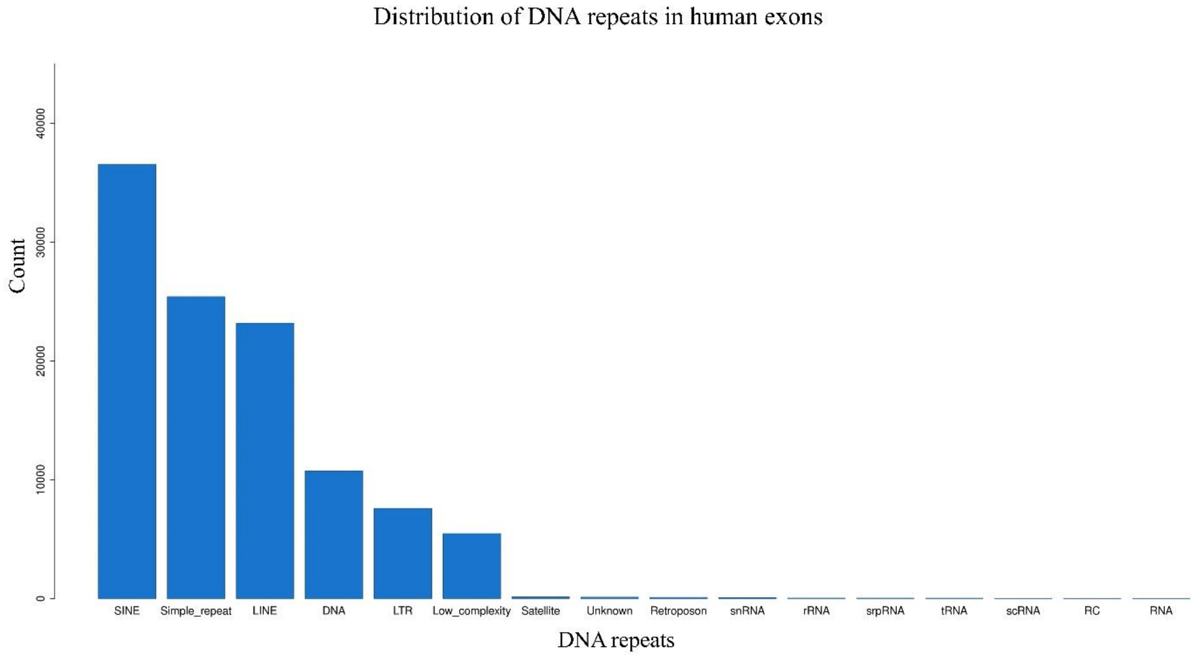


**Figure S1.** Distribution of DNA repeats in the human genome. The data about the DNA repeats is downloaded from UCSC genome browser, and have used the default classification criteria.

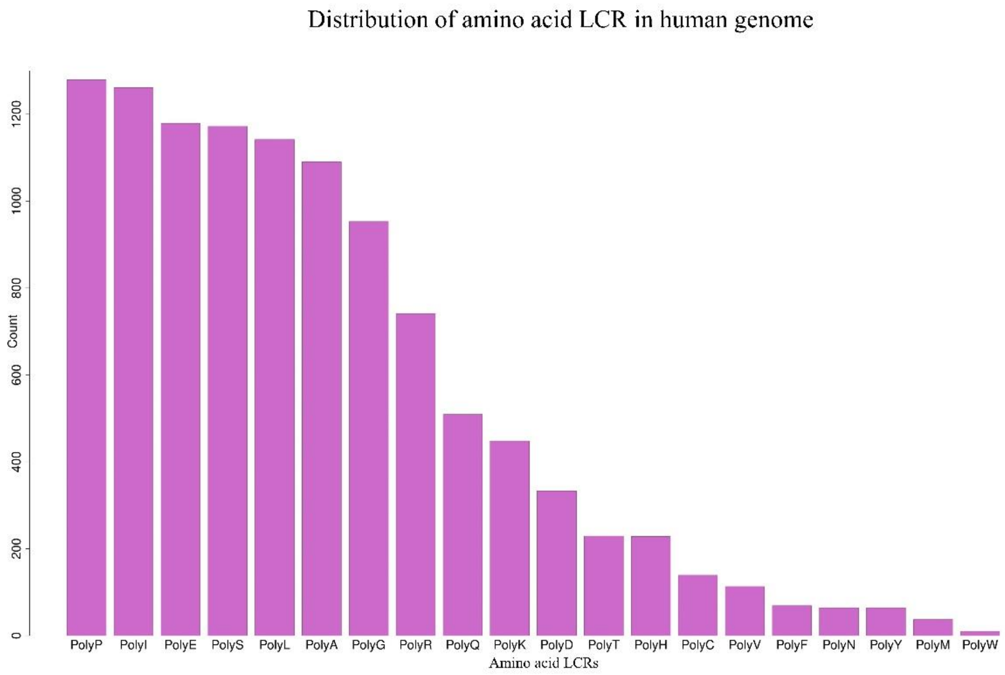


**Figure S2.** Distribution of amino acid LCR in the human genome.

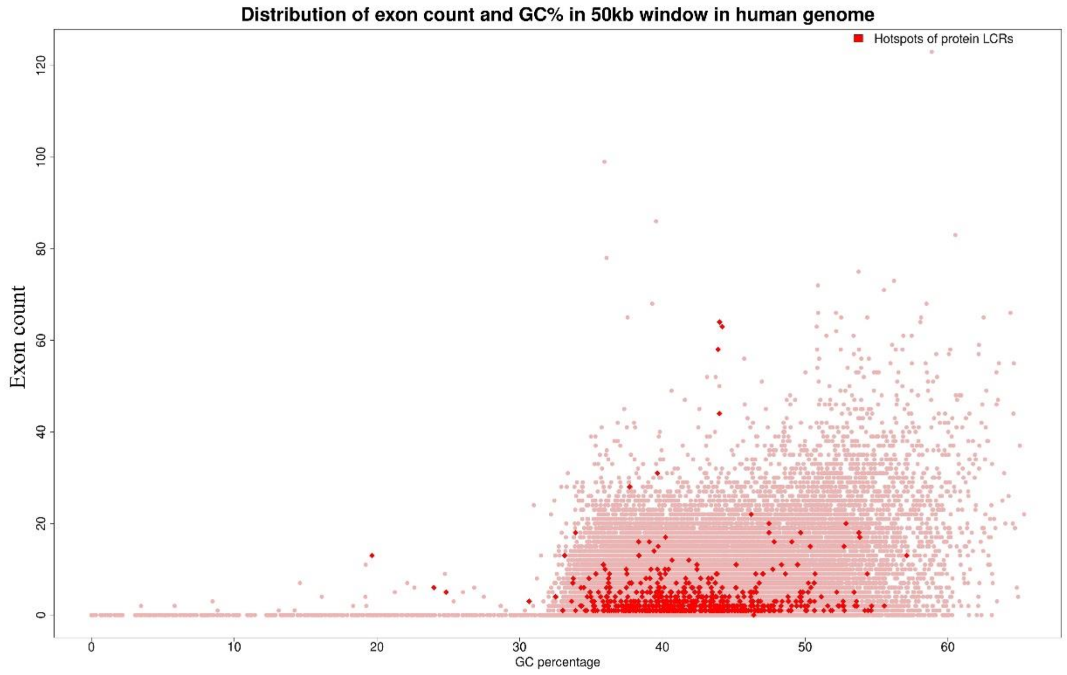


**Figure S3.** The scatter plot shows the distribution of exon count and GC% of LCR-containing exons along with other exons in the human genome.

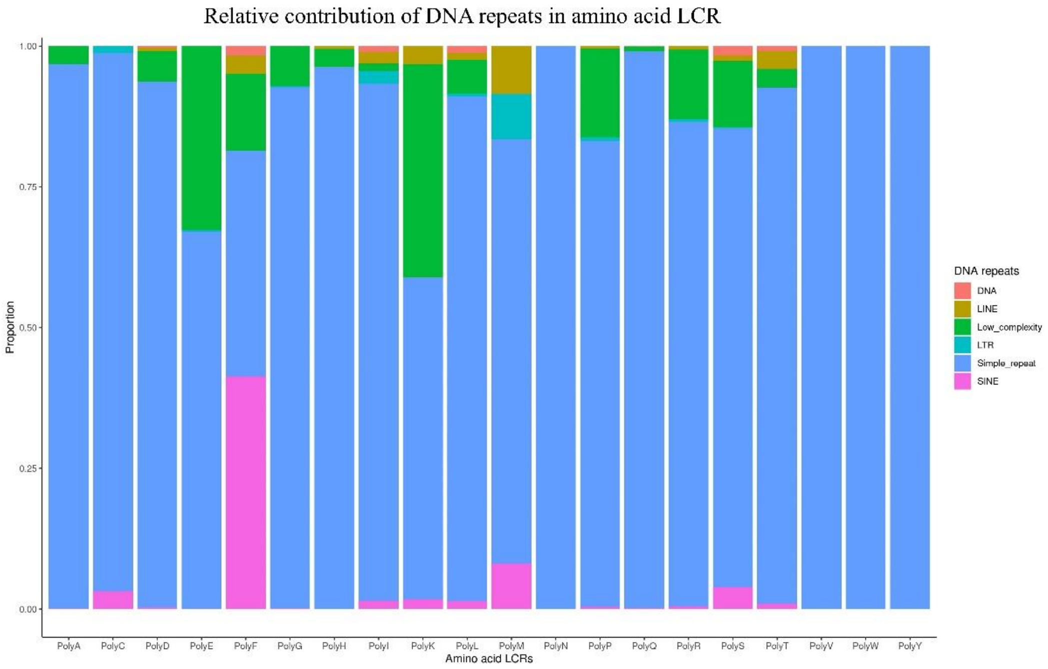


**Figure S4.** Relative contribution of different DNA repeats in the LCR in the human genome.

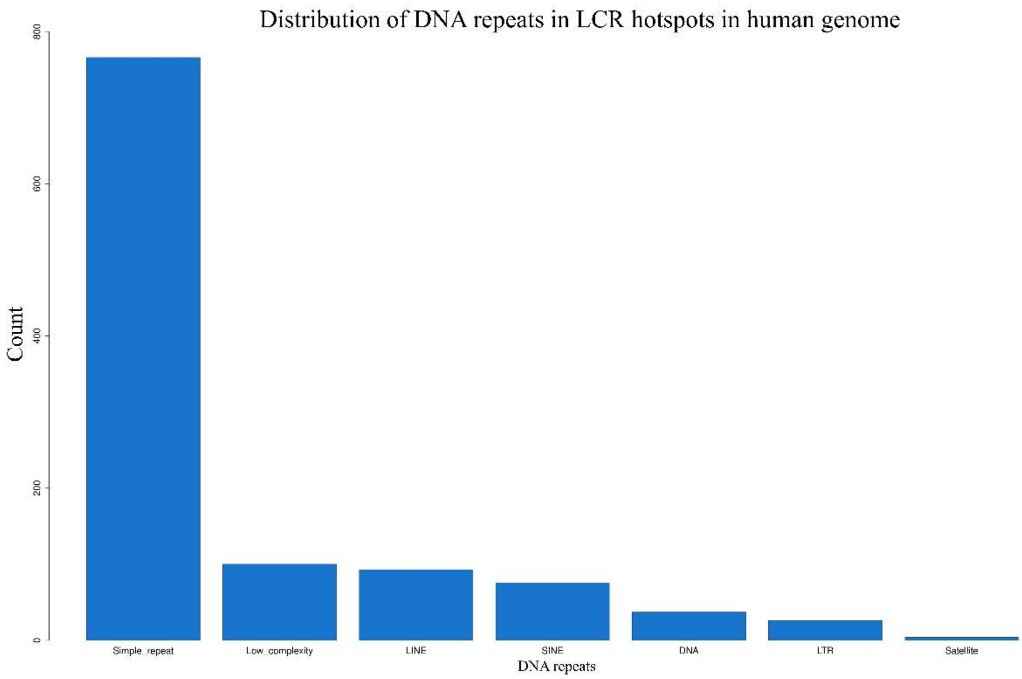


**Figure S5.** Simple DNA repeats are highest in count in the hotspots of LCR.

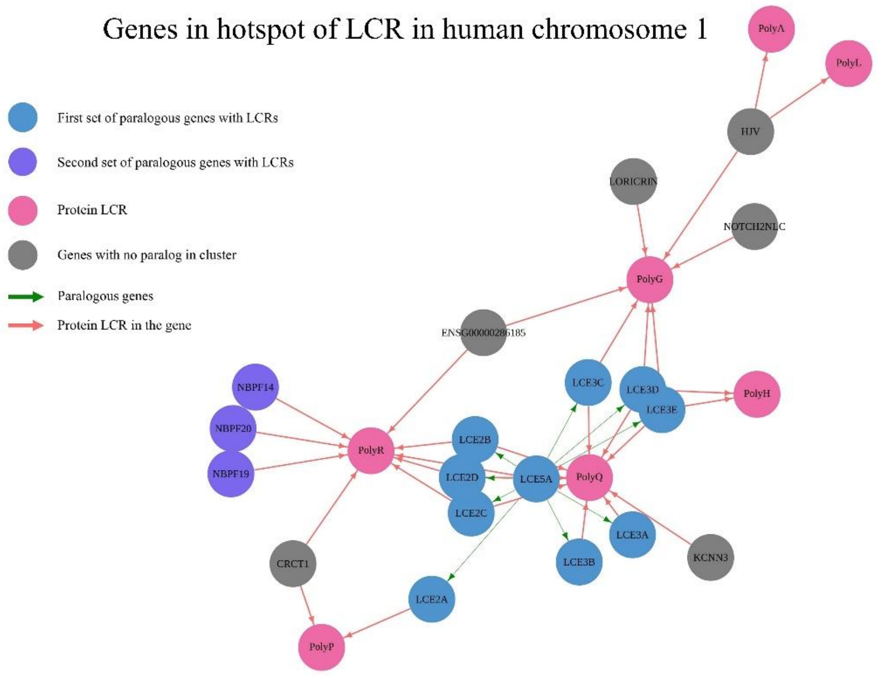


**Figure S6.** Cluster of LCR hotspot in chromosome 1 in the human genome. The paralogous genes are represented with the same vertex color while gray color represents absence of a paralogous gene in the cluster. Hotpink colored arrows represent the LCR shared between different genes, while green colored arrows represent paralogous genes.

**Figure S7.** Site frequency spectrum of minor allele frequency of LCR containing exons and exons without LCR in the human genome.

**Figure S8.** Major LCR involved in diseases. PolyA, polyG, polyL, and polyp are present in various hereditary diseases.

**Figure S9.** The network of LCR-containing disease-causing experimentally validated interacting genes. Distinct edge color represents distinct disease. The number of edges between two vertices represent the number of shared diseases between the genes. The pie chart within the vertices represents the proportion of distinct LCR within the gene.

**Figure S10.** The network of LCR-containing experimentally validated interacting genes in hereditary cancer-predisposing syndrome. The red color edges represent the hereditary cancer-predisposing syndrome. The pie chart within the vertices represents the proportion of distinct LCR within each gene.
