## Supplementary Figure 7 for "Patterns of low-complexity regions in human genes"

**Folded site frequency spectrum of amino acid LCR containing exons**

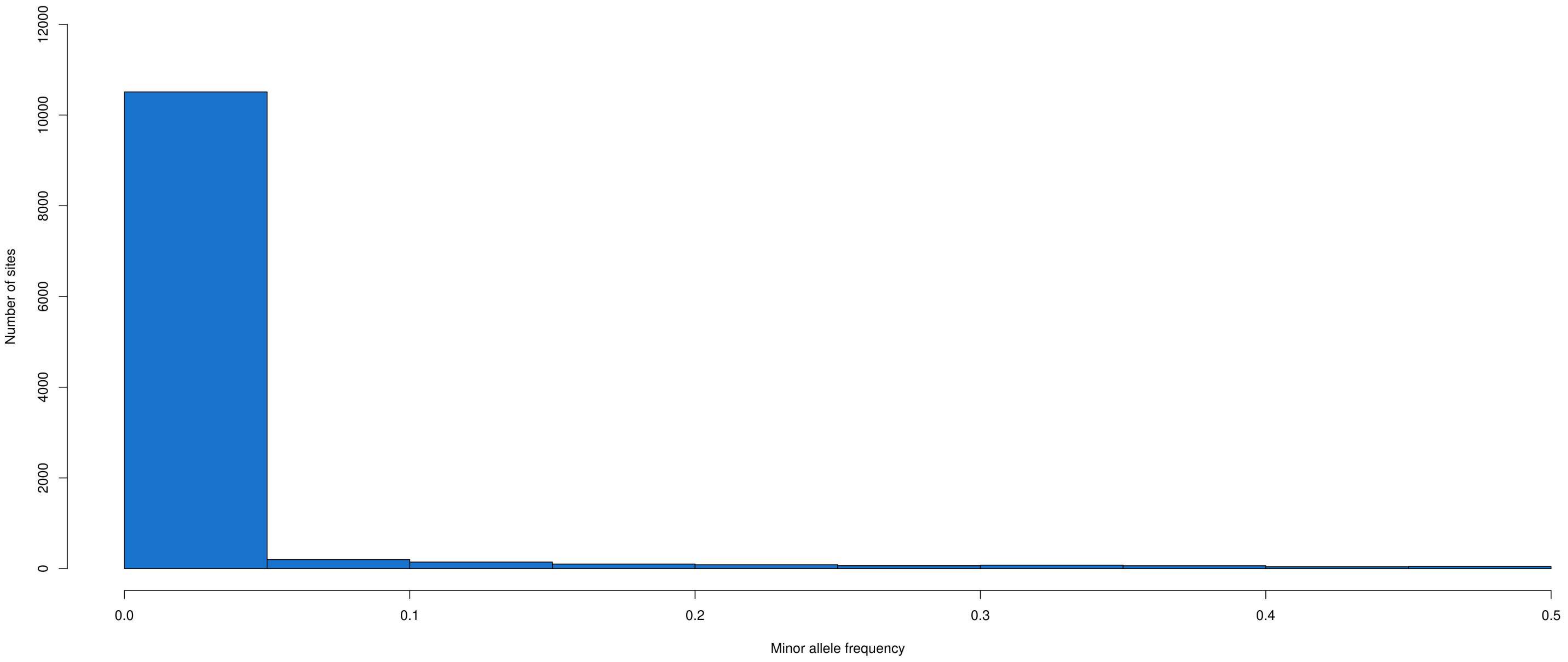


**Folded site frequency spectrum of exons without amino acid LCRs**

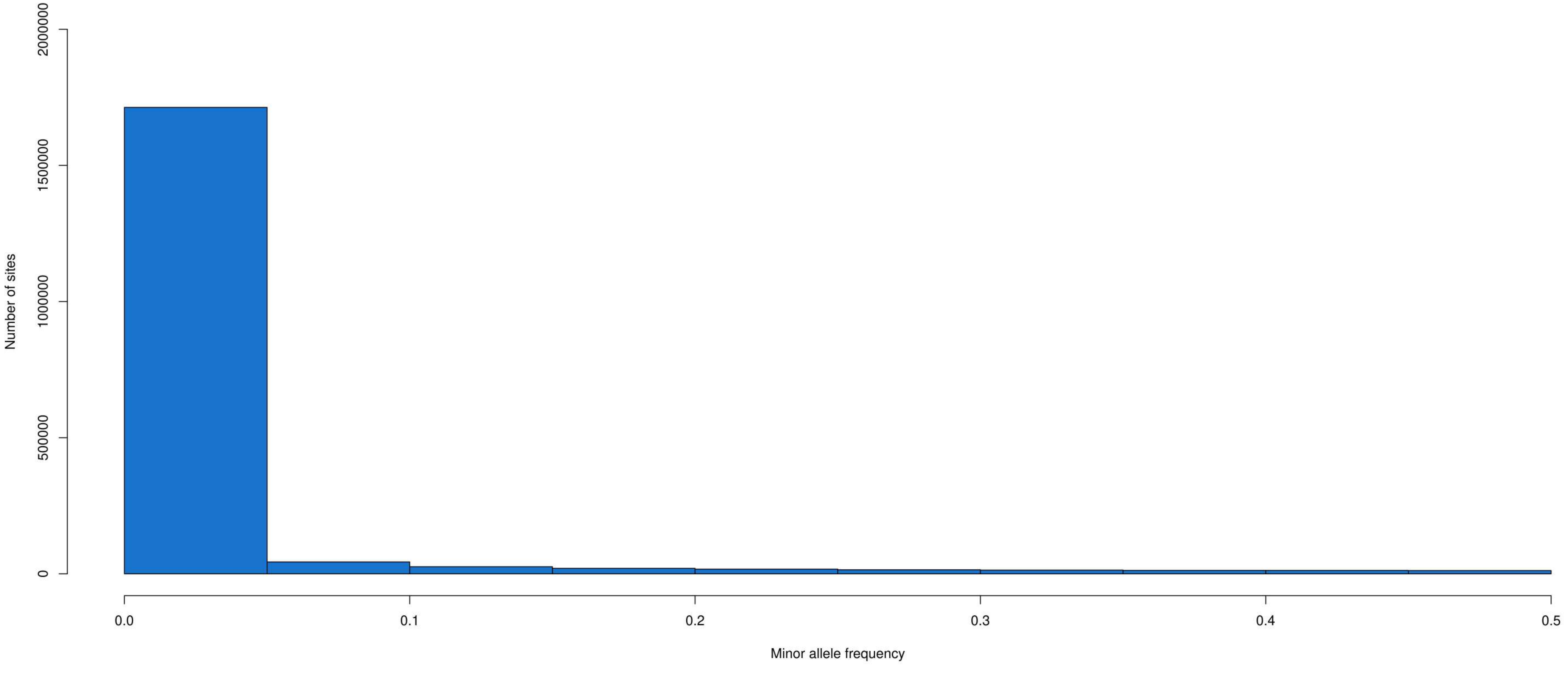
