## Supplementary figures and images for "Patterns of low-complexity regions in human genes"

### Supplementary Figure 8

#### Disorders due to LCRs

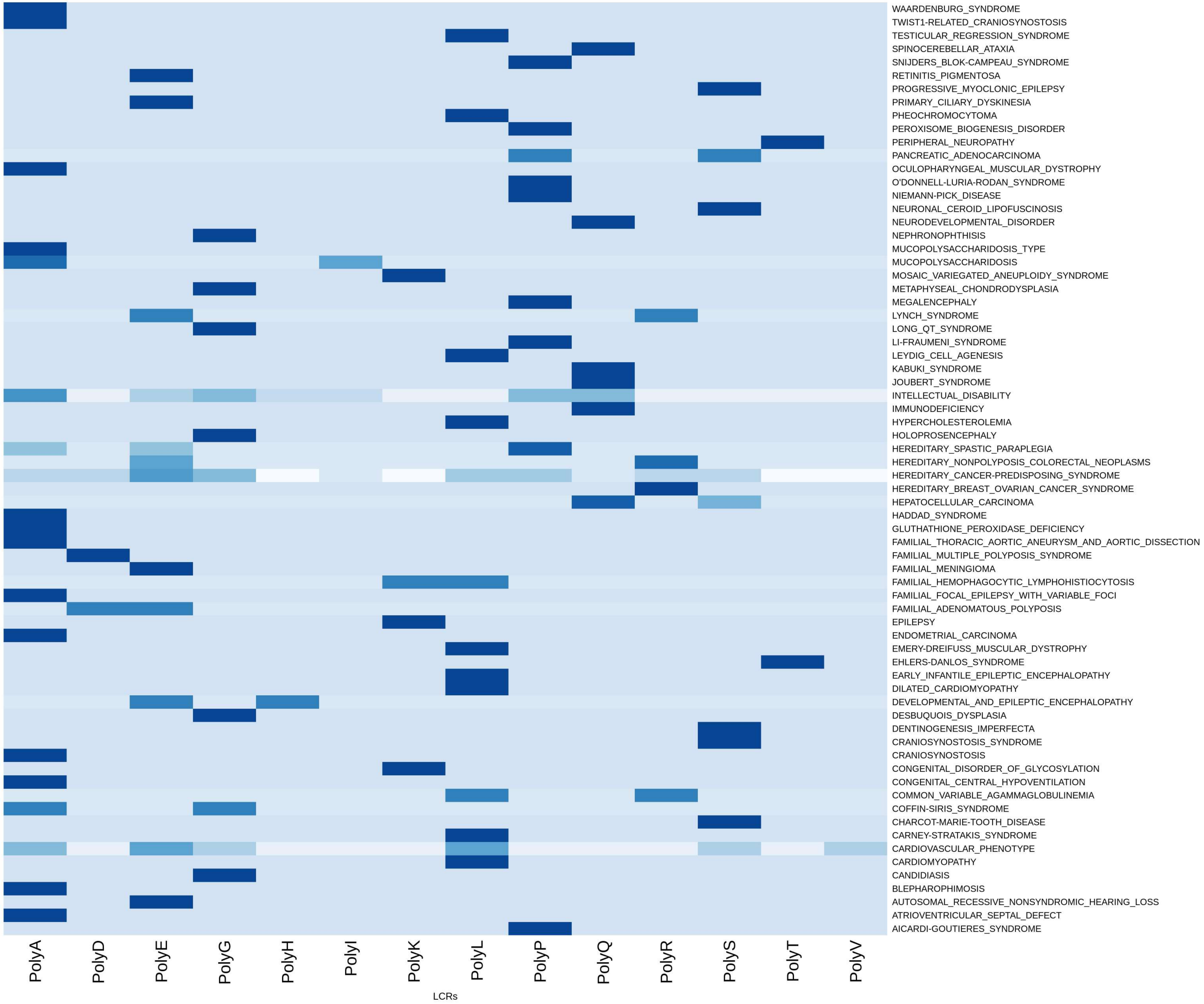

### Supplementary Figure 9

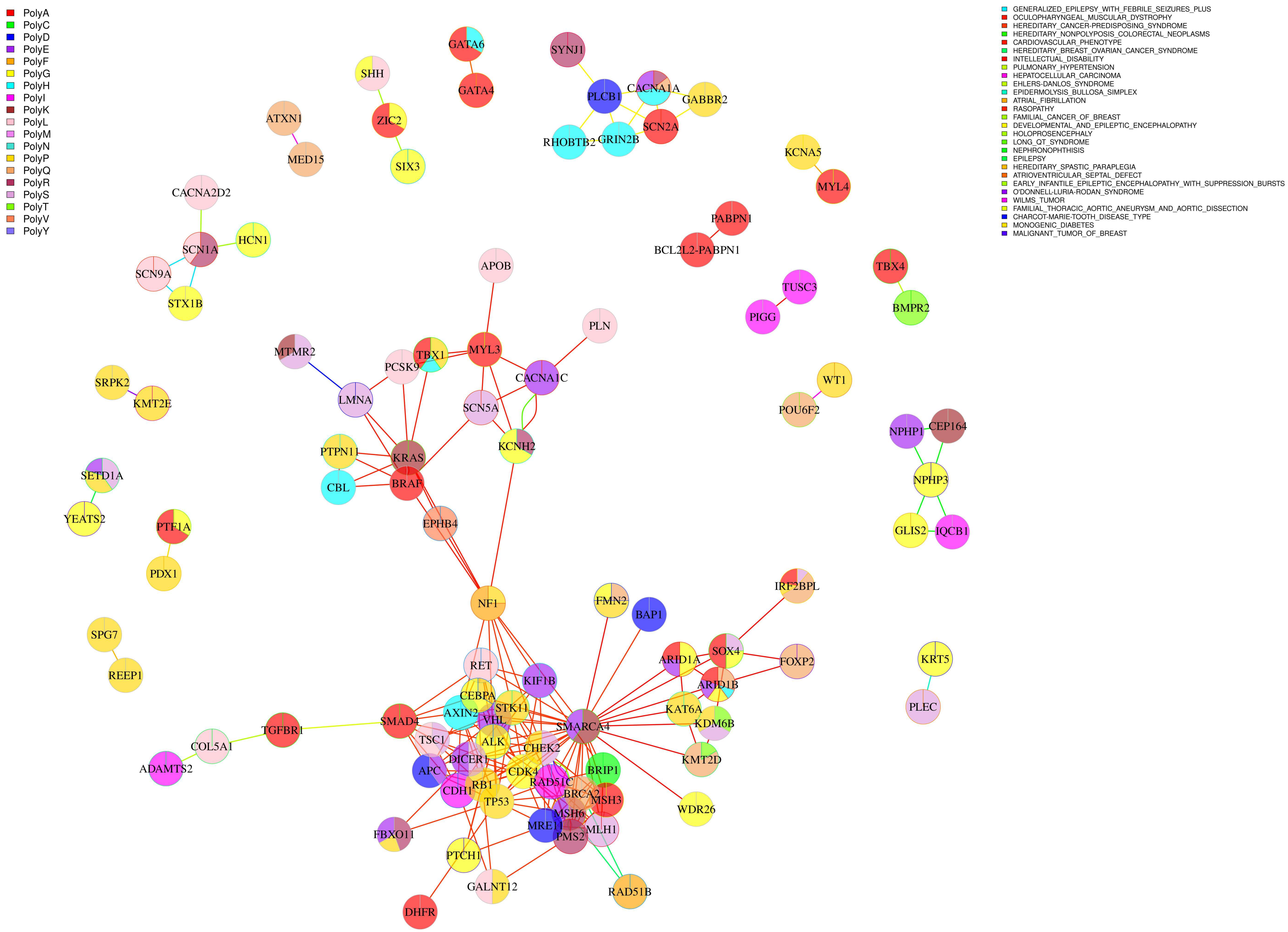

### Supplementary Figure 10

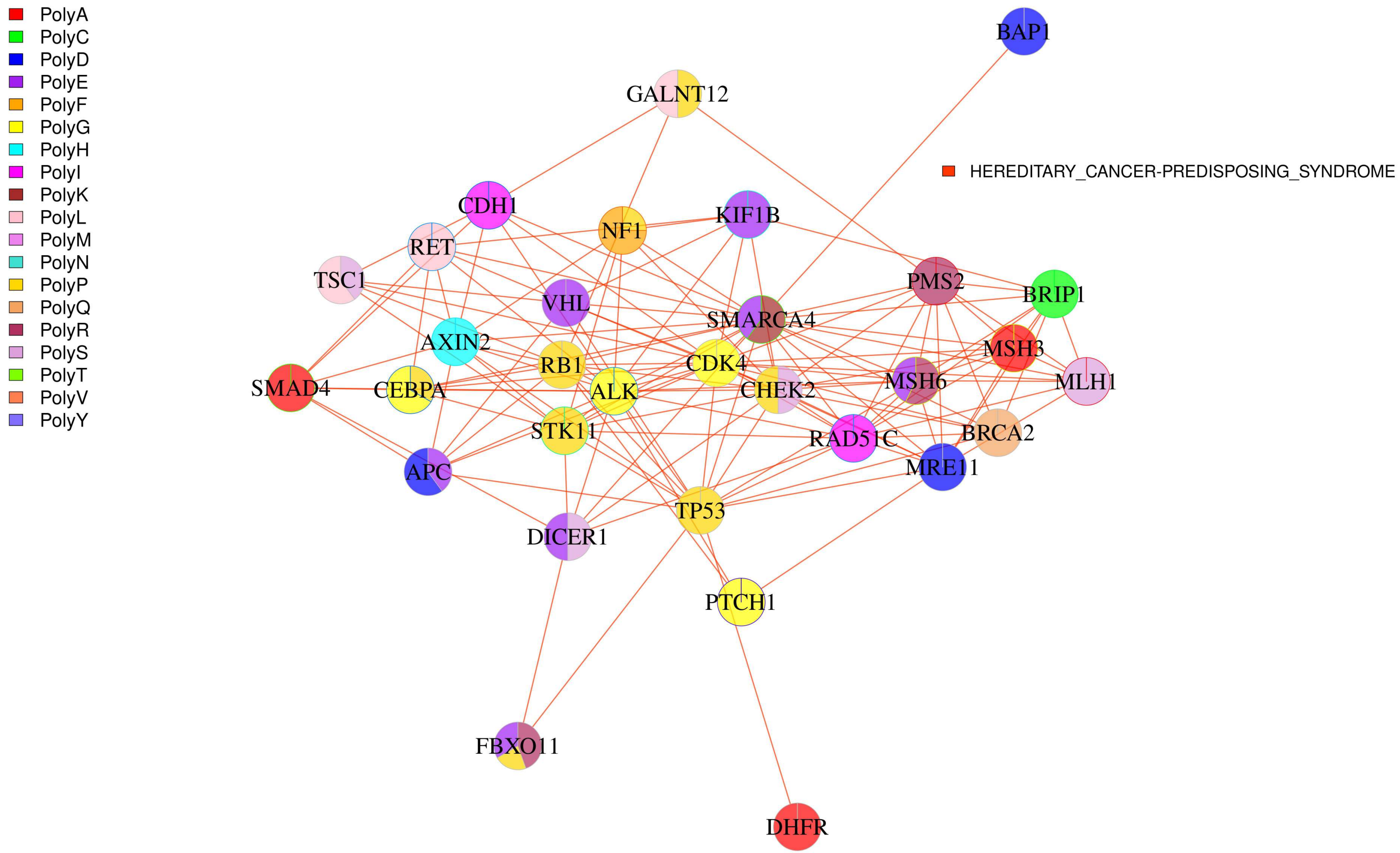
